# Ecological and anthropogenic drivers of large carnivore depredation on sheep in Europe

**DOI:** 10.1101/2020.04.14.041160

**Authors:** Vincenzo Gervasi, John D. C. Linnell, Tomaž Berce, Luigi Boitani, Rok Cerne, Benjamin Cretois, Paolo Ciucci, Christophe Duchamp, Adrienne Gastineau, Oksana Grente, Daniela Hilfiker, Djuro Huber, Yorgos Iliopoulos, Alexandros A. Karamanlidis, Francesca Marucco, Yorgos Mertzanis, Peep Männil, Harri Norberg, Nives Pagon, Luca Pedrotti, Pierre-Yves Quenette, Slaven Reljic, Valeria Salvatori, Tõnu Talvi, Manuela von Arx, Olivier Gimenez

**Affiliations:** CEFE, CNRS, University of Montpellier, University Paul Valéry Montpellier 3, EPHE, IRD, Montpellier, France; Norwegian Institute for Nature Research, PO Box 5685 Torgard, NO-7485 Trondheim, Norway; Slovenia Forest Service, Večna pot 2, 1000 Ljubljana, Slovenia; Dipartimento Biologia e Biotecnologie, Università di Roma Sapienza, Viale Università 32, 00185-Romae, Italy; Department of Geography, Norwegian University of Science and Technology, 7491 Trondheim, Norway; Dept. Biology and Biotechnologies, University of Rome La Sapienza, Viale dell’Università 32, 00185 Roma, Italy; Office National de la Chasse et de la Faune Sauvage, Gap, France; Equipe Ours, Unité Prédateurs Animaux Déprédateurs et Exotiques, Office Français de la Biodiversité, impasse de la Chapelle, 31800, Villeneuve-de-Rivière, France, Centre d’Ecologie et des Sciences de la Conservation (CESCO), Muséum National d’Histoire Naturelle, Centre National de la Recherche Scientifique, Sorbonne Université, CP 135, 43 rue Buffon, 75005, Paris, France; Unité Prédateurs Animaux Déprédateurs et Exotiques, Office Français de la Biodiversité, Micropolis - La Bérardie 05000 Gap, France. Centre d’Ecologie Fonctionnelle et Evolutive (CEFE), Centre National de la Recherche Scientifique, UMR 5175, Campus CNRS, 1919 Route de Mende, F-34293 Montpellier Cedex 5, France; Swiss Center for livestock protection, AGRIDEA, Eschikon 28, 8315 Lindau, Switzerland; Faculty of Veterinary Medicine, University of Zagreb, Heinzelova 55, 10000 Zagreb, Croatia; Callisto Wildlife and Nature Conservation Society, Greece; Arcturos – Civil Society for the Protection and Management of Wildlife and the Natural Environment, 53075 Aetos, Florina, Greece; University of Torino, Department of Life Sciences and Systems Biology, V. Accademia Albertina 13, 10123 Torino, Italy; Estonian Environment Agency, Mustamäe tee 33, Tartu, Estonia; Finnish Wildlife Agency, Rovaniemi, Finland; Parco Nazionale dello Stelvio, Glorenza (BZ), Italy; Equipe Ours, Unité Prédateurs-Animaux Déprédateurs, Office Français pour la Biodiversité, impasse de la Chapelle, 31800, Villeneuve-de-Rivière, France; Istituto di Ecologia Applicata - via B. Eustachio 10 - 00161, Rome, Italy; Environmental Board of the Estonian Ministry of Environment, Viidumäe, 93343 Saaremaa, Estonia; KORA – Carnivore Ecology and Wildlife Management, Thunstrasse 31, 3074 Muri b. Bern, Switzerland; Centre d’Ecologie Fonctionnelle et Evolutive - UMR 5175, Campus CNRS, 1919 Route de Mende, F-34293 Montpellier Cedex 5, France

**Keywords:** *Canis lupus*, carnivore conservation, compensation programs, *Gulo gulo*, human-wildlife conflict, impact, *Lynx lynx*, *Ursus arctos*

## Abstract

- Sharing space with large carnivores on a human-dominated continent like Europe results in multiple conflictful interactions with human interests, of which depredation on livestock is the most widespread. Wildlife management agencies maintain compensation programs for the damage caused by large carnivores, but the long-term effectiveness of such programs is often contested. Therefore, understanding the mechanisms driving large carnivore impact on human activities is necessary to identify key management actions to reduce it.
- We conducted an analysis of the impact by all four European large carnivores on sheep husbandry in 10 European countries, during the period 2010-2015. We ran a hierarchical Simultaneous Autoregressive model, to assess the influence of ecological and anthropogenic factors on the spatial and temporal patterns in the reported depredation levels across the continent.
- On average, about 35,000 sheep were compensated in the ten countries as killed by large carnivores annually, representing about 0.5% of the total sheep stock. Of them, 45% were recognized as killed by wolves, 24% by wolverines, 19% by lynx and 12% by bears. At the continental level, we found a positive relationship between wolf distribution and the number of compensated sheep, but not for the other three species. Impact levels were lower in the areas where large carnivore presence has been continuous compared to areas where they disappeared and recently returned. The model explained 62% of the variation in the number of compensated sheep per year in each administrative unit. Only 13% of the variation was related to the ecological components of the process.
- **Synthesis and Applications:** Large carnivore distribution and local abundance alone are poor predictors of large carnivore impact on livestock at the continental level. A few individuals can produce high damage, when the contribution of environmental, social and economic systems predisposes for it, whereas large populations can produce a limited impact when the same components of the system reduce the probability that depredations occur. Time seems to play in favour of a progressive reduction in the costs associated with coexistence, provided that the responsible agencies focus their attention both on compensation and co-adaptation.

## Introduction

The European continent is home to four species of large carnivores: brown bears (*Ursus arctos*), lynx (*Lynx lynx*), wolves (*Canis lupus*) and wolverines (*Gulo gulo*). After centuries of decline due to multiple causes (extermination policies, habitat destruction, reduction in the prey base, etc.) all these four species have progressively regained space, expanded their numbers, and recovered much of their former distribution during the last 50 years (Chapron et al., 2014). At present, 42 European large carnivore populations can be identified, 34 of which span over two or more (and up to nine) different countries (Chapron et al., 2014).

In the dichotomy between land sparing and land sharing conservation strategies (Phalan, Onial, Balmford, & Green, 2011), the European situation reveals that humans and large carnivores can share the same landscape, but not without a reciprocal impact. Due to the absence of large areas of wilderness in Europe (Venter et al., 2013), carnivores have almost entirely re-established their populations in rural, but highly human-modified landscapes, where humans raise livestock, keep bees for honey, hunt wild ungulates, and use forests and mountains for tourism and recreation (Chapron et al., 2014). Sharing space has therefore given rise to several forms of direct and indirect interaction between the ecological needs of large carnivores and the interests of rural humans (Bautista et al., 2019). These include depredation on livestock and destruction of beehives, dog killing, reduction of wild ungulate densities and other forms of impact that often generate conflicts which need to be managed (Linnell, 2013).

In response to large carnivore recovery, most European governments have introduced compensation programs for the damage they cause, as a way to increase social tolerance towards the species. Compensation programs rely on the social contract principle that the localized costs of human-large carnivore coexistence should be shared among all citizens (Schwerdtner & Gruber, 2007), under the expectation that time will allow the establishment of the appropriate coexistence mechanisms, thus progressively reducing the overall economic and social costs of the whole process. The long-term effectiveness of damage compensation programs in reducing large carnivore impact, though, is still under debate, considering that European countries nowadays pay almost 30 million euros per year for damage compensation, a sum that has increased during the last decade (Bautista et al., 2019). This raises the question if the whole compensation strategy will still be socially and economically sustainable in the near future (Linnell, 2013), especially considering that large carnivores will likely further expand their range in future years (Chapron et al., 2014). Therefore, understanding the mechanisms driving large carnivore impact on human activities is a necessary step, in order to identify those management actions which are more likely to reduce it.

Among the different forms of impact that large carnivore presence generates on human interests, depredation on livestock is by far the most widespread and relevant in economic terms (Linnell & Cretois, 2020). Livestock depredation is a very complex process, in which a large number of ecological and socio-economic factors interact at different spatial scales to determine the number of individuals encountered, killed, documented and compensated as large carnivore kills by the management authorities. Part of this process is just another type of predation, and therefore operates according to the same theoretical mechanisms of predation ecology (Linnell, Odden, & Mertens, 2012). The relative densities of large carnivores and their domestic prey, for instance, represent the numerical component of the depredation process in a classical sense, as formalized in the concepts of functional and numerical responses of predation (Holling, 1959). Therefore, the relative abundance of large carnivores with respect to their domestic prey is expected to affect the number of depredation events occurring each year in a given geographical context (Fig. 1). Additionally, landscape structure and land use can determine domestic prey encounter rates, accessibility and hunting success by large carnivores, similarly to the way they can modulate predation risk and kill rates in the wild (Kauffman, Smith, Stahler, & Daniel, 2007). Finally, the availability of alternative wild ungulate prey can affect the tendency by large carnivores to rely on domestic species, similarly to the way prey selection patterns and predatory behaviour are influenced by relative prey densities in multi-prey systems in the wild (Ciucci et al. 2020).

**Fig. 1.**
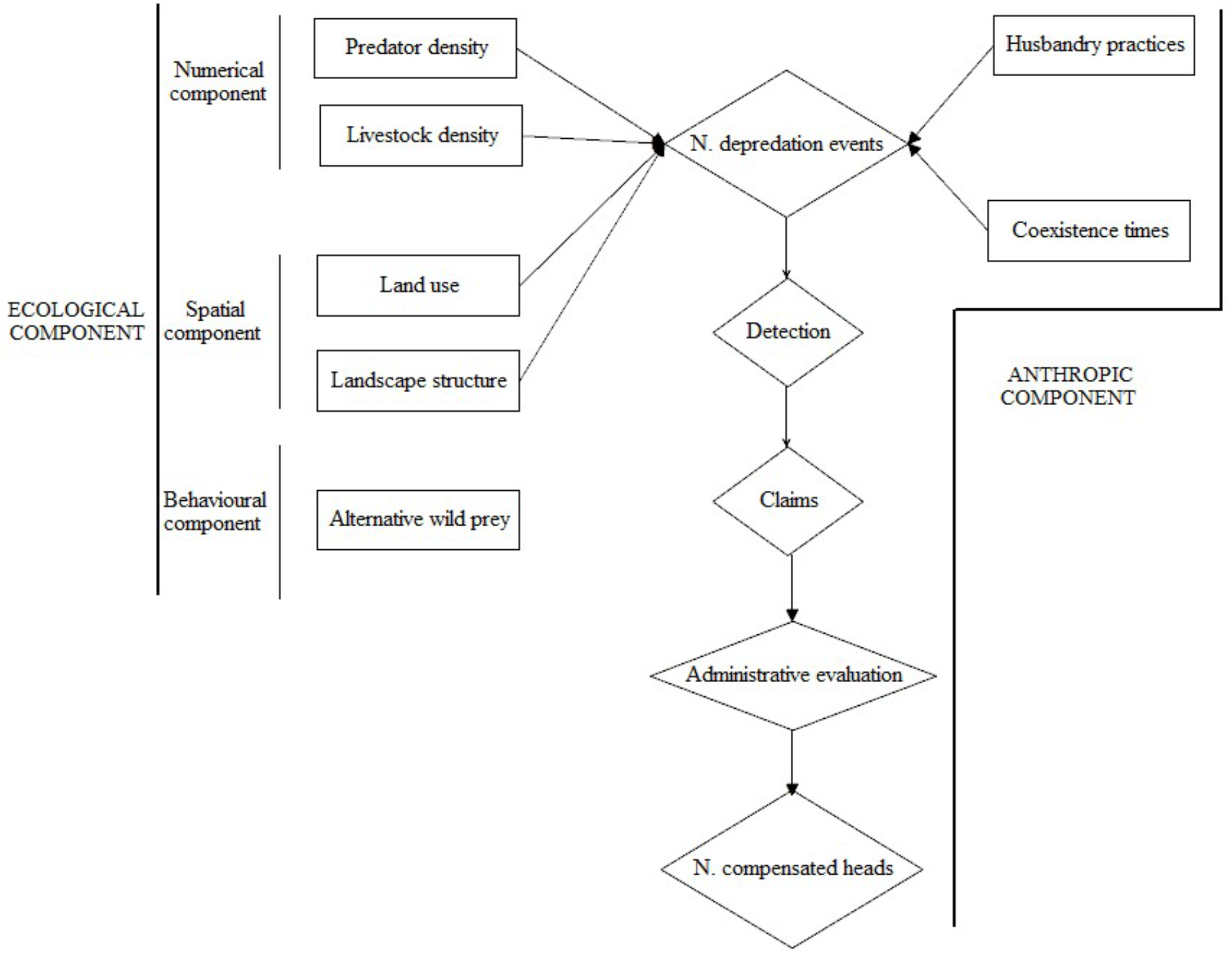
Conceptual diagram of the ecological and anthropogenic mechanisms generating the number of annually compensated sheep losses to large carnivores. The diagram also illustrates the analytical framework used to analyse the spatial and temporal variation in the number of compensated sheep head in 10 European countries, years 2010-2015.

The main challenge in the study of large carnivore depredation on livestock, though, is that the purely ecological mechanisms (density, habitat structure, predator behaviour) are only one component of the process, and possibly not the most relevant in determining its magnitude and spatial variation. Cultural, historical, economic and social aspects of the interaction between humans, livestock and large carnivores are crucial in affecting the long causal chain that determines the costs of coexistence. For instance, livestock husbandry practices, which are highly influenced by local historical and cultural traits, can strongly affect predation risk and the resulting magnitude of the depredation process (Eklund, López-Bao, Tourani, Chapron, & Frank, 2017). They can also change and progressively adapt to the need of reducing depredation risk, thus generating the expectation that longer periods of co-occurrence will allow the establishment of the appropriate mutual adaptation mechanisms, especially if supported by effective management actions (Carter & Linnell, 2016). Additionally, in most of the cases depredation events are neither directly nor accurately observed. Instead, they derive from a long chain of events that starts when the actual depredation occurs, implies a certain probability to detect the event, continues with a farmer’s willingness to report it and claim compensation, and includes a different set of evaluation methods by local management authorities. Such process ends with an administrative decision to classify the event as a depredation, and therefore refund the farmer (see the diagram in Fig. 1 for an illustration of the ecological and anthropogenic factors linking predation ecology, livestock depredations and compensated losses). Therefore, looking at depredation through the filter of the different compensation systems requires accounting for the risk of getting a biased image of its relative magnitude in the different contexts. Although the dual nature of livestock depredation as both an ecological and a socio-economic process is a well-established concept (Linnell, 2013), a formal evaluation of their relative importance in affecting the spatial and temporal variation in depredation and compensation patterns has not yet been performed.

Building on the above-described conceptual framework, we analysed the impact of all four European large carnivores on sheep husbandry in 10 European countries, during the period 2010–2015. We collected data about the prevalent husbandry practices, the characteristics of the compensation schemes and the number of confirmed depredation events in each of the administrative units in charge of large carnivore compensation in each country. Then, we ran a hierarchical Simultaneous Autoregressive model (SAR), to assess the influence of some ecological and anthropogenic factors on the emerging spatial and temporal patterns in depredation levels across the continent. We focused on sheep depredation, as sheep alone represent more than 60% of all the compensation payments in Europe (Linnell & Cretois, 2020), thus being the most relevant form of material impact of large carnivores on human interests, from an economic point of view.

In particular, we focused on the following research hypotheses:

1. The area occupied by large carnivores in a given area is a predictor of the number of verified sheep depredations;
2. There are differences among the four large carnivore species, in terms of their relative impact on livestock husbandry;
3. The geographic variation in land use, habitat types and landscape structure affects the spatial variation in compensation patterns among European countries;
4. Recently re-colonized areas are more impacted by large carnivores than the ones in which humans and large carnivores share a longer history of co-occurrence;
5. A higher number of alternative wild ungulate species available corresponds to a reduction in large carnivore impact on sheep in a given area;
6. The ecological component of the depredation process (numerical, spatial, behavioural) is the most relevant in influencing the magnitude of large carnivore impact on livestock.

## Methods

### Data collection

We obtained data from 10 European countries, namely Croatia, Estonia, Finland, France, Greece, Italy, Norway, Slovenia, Sweden and Switzerland. Data from Italy were limited to the Alpine wolf and bear populations (Chapron et al., 2014). We chose the above-mentioned countries and regions because they allowed us to cover a north-south geographical gradient of the European continent, which involved a set of environmental, social, and economic differences. The choice was also based on the availability of organised and accessible national or regional datasets, which contained the type of information needed to compile the review and run the subsequent analyses. We collected data according to the NUTS3 (Nomenclature of Territorial Units for Statistics) classification, which corresponded in most countries to the administrative level of departments, cantons, provinces, etc.

For each year and each NUTS3 unit, we collected data about the estimated abundance of each large carnivore species whenever available, or the minimum number of individuals known to be present. We also collected the number of registered sheep and the number of sheep compensated as killed by large carnivores. Additionally, for each country, we compiled a summary description of the prevalent sheep husbandry practices, of the most common damage prevention systems employed by sheep farmers, and of the main characteristics of the national compensation system, whose results are summarized in Table S4 and in the Appendix 1 in the Additional Supporting Information. We received data from national and regional wildlife agencies, from published literature and reports, as well as from researchers and practitioners. The complete description of the data sources for each data type included in the review is available in tables S1, S2 and S3.

### Modelling

To explore the main patterns in the number of sheep heads compensated each year as killed by large carnivores in the 10 countries included in the study, we used a Bayesian hierarchical SAR Poisson models (Zhu, Zheng, Carroll, & Aukema, 2008) in Jags (Plummer, 2003). One of the objectives of our study was to test and estimate the effect of large carnivore abundance on the expected number of annually-compensated sheep (hypothesis 1). As not all countries included in the study were able to provide large carnivore abundance data at the NUTS3 spatial resolution, the surface of the species distribution area in each sampling unit was the only common metric we could resort to. The relationship between the area occupied by a species and the number of individuals living in that area, though, is not expected to be a constant (Carbone & Gittleman, 2002). Habitat productivity, body size and several other factors influence home range size and the area needed to sustain a given animal population (Harestad & Bunnel, 1979; Nilsen, Herfindal, & Linnell, 2005). Therefore, the use of distribution as a proxy for abundance, at the scale of the whole European continent, could potentially introduce a bias in all subsequent analyses. In order to account for and prevent such bias, we built the first level of the hierarchical SAR Poisson model (Eq. 1) to analyse the species-specific area/abundance relationship for each of the four large carnivore species. To this aim, we defined the number of individuals of each large carnivore species *s* detected in each NUTS3 region *i* on year *t* (period 2010-2015) as a Poisson random variable with parameter (γ*_s,i,t_*). This parameter was modelled (on the log scale) as a function of the area occupied by the species in the same region. To account for the large-scale spatial variation in climate and habitat productivity, we included the latitude of each NUTS3 region in the model as a predictor. As large carnivore home range size is also influenced by prey availability, we used presence/absence distribution maps for the main wild ungulate species in Europe (roe deer, red deer, wild boar, moose, chamois, wild reindeer; Linnell & Cretois, 2020) and calculated the number of wild ungulate prey species available in each NUTS3 unit. We used this factor variable as an additional predictor for large carnivore abundance. To account for the spatial correlation of neighbouring NUTS3 units, we also added a normally distributed individual random term *ε_i,s_* for each region *i* and species *s* in the model. The random effect had mean equal to zero and variance defined as σ^2^ (*D – φW*), in which *σ* was the standard deviation, *W*was a binary adjacency matrix (1 = bordering, 0 = not bordering), *D* was the diagonal matrix of *W*, and *φ* was an estimated parameter controlling the intensity of the spatial correlation. Finally, we also added a time-dependent random effect *τ_t,s_* accounting for the nested structure of the data, in which six abundance data points (one for each year) were available for each large carnivore species in each region. A log link function was used to run the Poisson regression model.

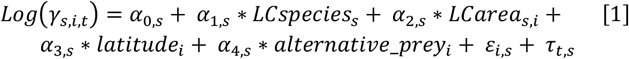

The second level of the Bayesian hierarchical model (Eq. 2) was meant to interpret part of the variation in the number of compensated sheep heads in each NUTS3 unit and in each country. Model structure was similar to the one used for the first level of the model. We initially ran the model using a common intercept and slope for all the four large carnivore species, in order to reveal any common pattern in compensation levels. Then, we ran another version of the model, which included a separate intercepts and slopes for each large carnivore species, with the aim to highlight species-specific patterns and the relative impact of each large carnivore species (hypothesis 2). We used sheep abundance and the index of large carnivore abundance (derived from Eq. 1) as linear predictors, in order to include the numerical component of the depredation process and to test to what extent the area occupied by large carnivores in each NUTS3 unit affected the resulting number of compensated losses. We also included three macroscopic spatial variables, to test for the effect of land use and landscape structure on the sheep compensation process (hypothesis 3). Using a Digital Elevation Model for Europe (DEM, resolution 25 meters) and the Corine Land Cover map (EEA-ETC/TE, 2002), we extracted the proportion of land occupied by forest (conifer, broadleaved or mixed), the edge density index as an estimate of the availability of ecotone areas, and the landscape ruggedness index for each NUTS3 spatial unit. We added these variables as three additional linear predictors in the Poisson regression model (Eq. 2). To test for the effect of time since large carnivore re-colonization (hypothesis 4), we overlaid the study area with the estimated large carnivore distribution referring to the period 1950-1970 (Chapron et al., 2014), and produced a binary variable for each NUTS3 region, indicating if a given large carnivore species was already present at that time or returned in more recent years. Similarly to what was done for the first level of the hierarchical model, we used the number of wild ungulate prey available in each sampling unit as an additional predictor of compensation levels, under the hypothesis that a wider spectrum of alternative wild prey would reduce the number of compensated sheep heads (hypothesis 5). Three additional random effects were added to the depredation model: an individual random effect *μ_i,s_* for each region *i* and species *s*, accounting for the spatial auto-correlation in the data in the same way as described for the first level of the hierarchical model; a time-specific random effect *θ_t,s_* for each year *t* and species *s*; a country and species-specific random effect *ρ_k,s_*, which estimated the residual variation in compensated sheep heads, which could not be explained by the other terms of the model. With respect to the conceptual differentiation between ecological and anthropogenic predictors of large carnivore damage, the explicit variables represented the ecological component of the process (numerical, spatial, behavioural), whereas the effect of the anthropogenic factors was summarized through the random effects.

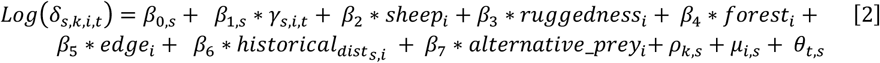

Finally, in order to separate the effects of the ecological (explicit) and anthropogenic (implicit) factors in affecting the compensation process, we also predicted the number of compensated sheep heads using a model which excluded the individual and country-specific random effects. This allowed us to produce an estimate of what compensation levels would be expected in a country, if only the numerical, spatial and behavioural component of the depredation process were in action. The comparison of these predictions with the observed compensation levels allowed us to infer the positive/negative effect of the additional country-specific components that were not explicitly tested in the depredation model. We also estimated the proportion of variance explained by the two models (R^2^), in order to highlight the relative importance of the explicit and implicit terms in the compensation process (hypothesis 6). To this aim, we calculated the difference between the model residuals and the residuals of an intercept-only model (Nakagawa & Schielzeth, 2013). We used a log link to run also this part of the Poisson model. Models converged in Jags, using 10,000 iterations and a burning phase of 5,000 iterations.

## Results

Overall, the 10 countries considered in the analysis hosted about 26 million sheep, of which about 7.6 million (29%) overlapped with the distribution of at least one large carnivore species (Tab. 1). In the same geographic area, a minimum of about 2,000 wolves, 7,600 bears, 1,300 wolverines and 5,600 lynx were estimated to live (Tab. 1), for a total of 16,500 individuals. On average, about 35,000 sheep were annually compensated in the ten countries as killed by large carnivores (Tab. 1 and Fig. 2). Out of them, about 45% were recognized as killed by wolves, 12% by bears, 24% by wolverines and 19% by lynx. In average, 7.7 sheep were compensated for each wolf at the continental level, 0.55 sheep for each bear, 6.55 for each wolverine and 1.17 for each lynx.

**Fig. 2.**
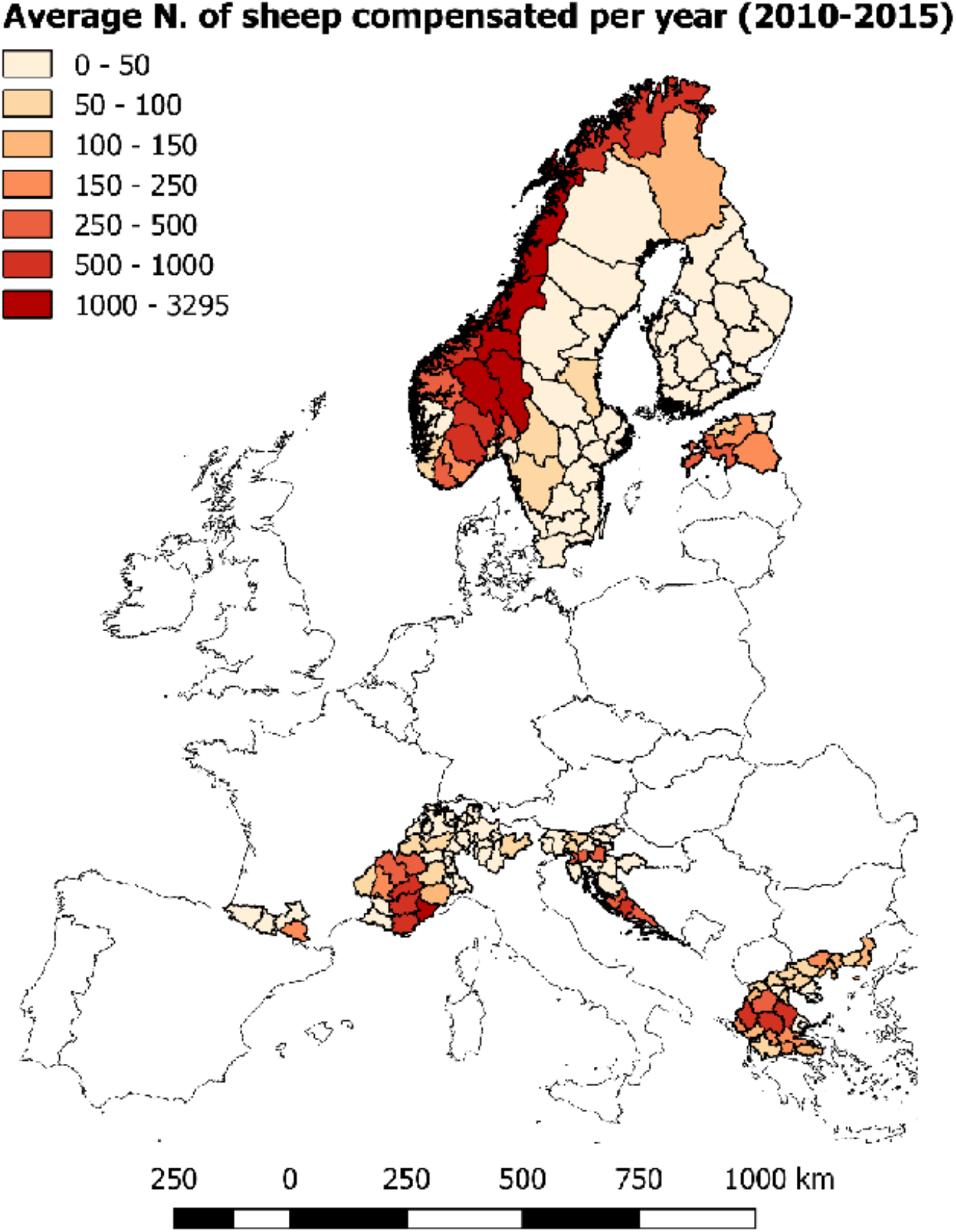
Average number of sheep heads totally compensated as killed by large carnivores in 171 administrative units and 10 countries in Europe (NUTS3 level).

**Tab. 1.** Summary statistics of sheep husbandry, large carnivore estimated abundance and total compensated sheep heads in the 10 European countries included in the large carnivore impact analysis, years 2010-2015.

| Country | Sheep abundance in LC | Large carnivore abundance |  |  |  | N. compensated heads per year (mean) |  |  |  |  |
| --- | --- | --- | --- | --- | --- | --- | --- | --- | --- | --- |
|  | distribution areas | (Minimum number detected) |  |  |  |  |  |  |  |  |
|  | (thousands) | Wolf | Bear | Wolverine | Lynx | Wolf | Bear | Wolverine | Lynx | Total |
| Croatia | 418 | 193 | 1000 | 0 | 50 | 1674 | 1 | 0 | 0 | 1675 |
| Estonia | 91 | 230 | 650 | 0 | 460 | 806 | 5 | 0 | 23 | 834 |
| Finland | 134 | 157 | 1700 | 240 | 2485 | 85 | 164 | 0 | 32 | 281 |
| France | 998 | 250 | 25 | 0 | 108 | 5285 | 289 | 0 | 0 | 5574 |
| Greece | 4729 | 700 | 450 | 0 | 0 | 3972 | 229 | 0 | 0 | 4201 |
| Italy (Alps) | 217 | 157 | 35 | 0 | 0 | 251 | 117 | 0 | 0 | 368 |
| Norway | 330 | 33 | 105 | 360 | 396 | 2037 | 2942 | 8469 | 6095 | 19543 |
| Slovenia | 81 | 46 | 608 | 0 | 20 | 1083 | 478 | 0 | 6 | 1567 |
| Sweden | 489 | 295 | 3300 | 692 | 1650 | 308 | 23 | 0 | 463 | 794 |
| Switzerland | 224 | 13 | 0 | 0 | 166 | 220 | 0 | 0 | 16 | 236 |
| Total | 7711 | 2074 | 7873 | 1292 | 5335 | 15721 | 4248 | 8469 | 6635 | 35073 |

In absolute terms, Norway was the country with the highest number of compensated sheep heads (N = 19,543, 54% of the total, see Tab. 1) followed by France (N = 5,574) and Greece (N = 4,201). Finland, Sweden and Switzerland exhibited the lowest absolute numbers of compensated heads, with an average of less than 1,000 compensated heads per year (Tab. 1). In relative terms, Norway was still the country suffering the highest costs of sheep-large carnivore coexistence, as about 5.6% of all sheep living in the country were compensated as killed by one of the four large carnivore species each year. All the other countries lost less than 1% of their national sheep flock to large carnivores.

### Drivers of damage compensation across Europe

For all the four large carnivore species, the first level of the Bayesian hierarchical model highlighted a significant and positive relationship between the area occupied by the species in each NUTS3 unit and the number of individuals detected by the monitoring system. Species-specific slopes for this relationship varied between 0.048 for lynx (SD = 0.015, 95% CIs = 0.019 – 0.079) and 0.327 for wolves (SD = 0.074, 95% CIs = 0.181 – 0.470). The effect of latitude on the area/abundance relationship was only significant for wolves (β = −0.069, SD = 0.030, 95% CIs = −0.139 – −0.019), but not for the other three species. At the average latitude, 549 km^2^ of permanent distribution area were needed to host one wolf territory (Fig. 3a). This value increased to 1,369 km^2^ at the northernmost latitude and decreased to 216 km^2^ at the southernmost latitude.

**Fig. 3.**
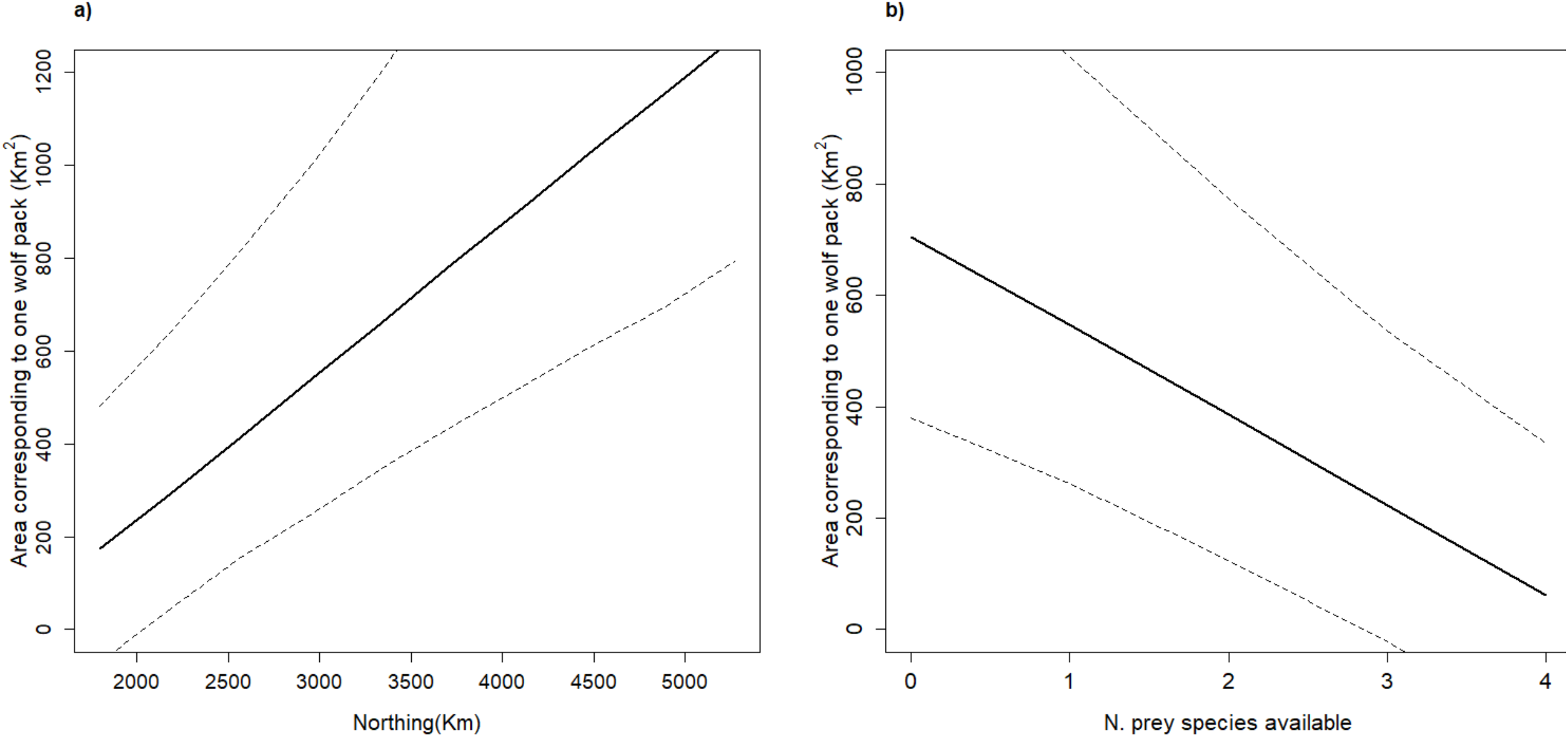
Relationship between latitude (a), the number of wild ungulate prey species available (b) and the area corresponding to one wolf territory in Europe.

The model also revealed a significant effect of the number of wild ungulate species available in a given area on the area/abundance relationship for wolves (β = 0.498, SD = 0.149, 95% CIs = 0.219 – 0.788) and lynx (β = 0.933, SD = 0.287, 95% CIs = 0.406 – 1.485). As shown in Fig 3b for wolves, the higher was the number of available wild prey species, the smaller was the distribution area required for one wolf territory. Overall, the first level of the hierarchical model revealed that the use of large carnivore distribution area, corrected by the above-mentioned factors, was a reliable proxy for large carnivore abundance in each NUTS3 unit.

The second level of the Bayesian hierarchical model revealed a significant positive relationship between the area occupied by large carnivores in each NUTS3 administrative unit and the number of compensated sheep (hypothesis 1; β = 0.012, SD = 0.001, 95% CIs = 0.011-0.013). A significant positive relationship also existed between sheep abundance and the number of sheep compensated (β = 0.084, SD = 0.029, 95% CIs = 0.024-0.141). Both these slopes refer to a model comprising a pooled effect for all the four large carnivore species considered in the analysis. When parameterizing the model with species-specific intercepts and slopes, the model revealed significant differences between the four large carnivore species (hypothesis 2). After accounting for all the other factors, verified wolf damage was significantly higher than that attributed to the other three species, as indicated by the higher intercept value in the model. In addition, wolves were the only species exhibiting a significant positive relationship between their distribution area and the expected number of compensated sheep per year (β = 0.131, SD = 0.004, 95% CIs = 0.123-0.139). The model reported no significant effects of any of the landscape variables (hypothesis 3), but it did reveal a significant effect of the historical continuity of large carnivore presence in reducing the expected number of compensated sheep per year (hypothesis 4; β = −0.973, SD = 0.471, 95% CIs = −1.914 – −0.069). The number of alternative wild ungulate prey species available in a given geographic area did not correspond to a reduction in the expected large carnivore impact on sheep husbandry (hypothesis 5; β = −0.042, SD = 0.247, 95% CIs = −0.516 – 0.449).

The estimation of random effects in the second level of the hierarchical model revealed large differences in the expected compensation levels among countries and among large carnivore species, a pattern that was also confirmed by the comparison between the observed number of sheep annually compensated and the one predicted by a model which accounted only for the ecological component of the process (Fig. 4). Norway, for example, was predicted to generate 4,348 compensated sheep per year, as opposed to the 19,543 actually observed. Similarly, France reported more than 5,000 compensated heads per year, while the explicit part of the model predicted no more than 400. On the other hand, Sweden and Finland generated only 10-15% of the damage levels predicted by the number of large carnivores present in those countries and by the size of their national flocks (Fig. 4).

**Fig. 4.**
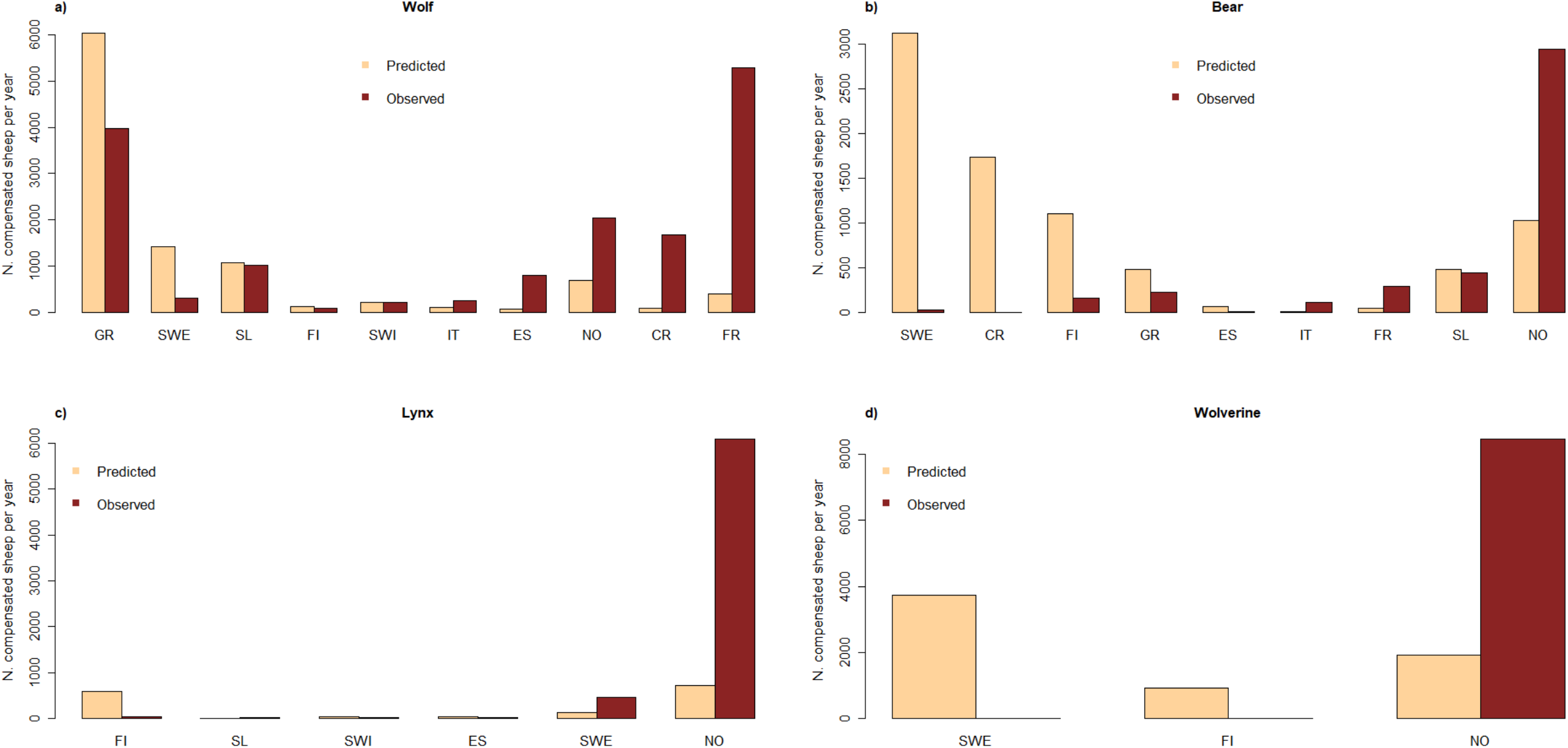
Comparison between the observed sheep compensation frequencies referring to four large carnivore species in 10 European countries and the ones predicted by the Bayesian hierarchical Simultaneous Autoregressive model (CR = Croatia; ES = Estonia; FI = Finland; FR = France; GR = Greece; IT = Italy (Alps); NO = Norway; SL = Slovenia; SWE = Sweden; SWI = Switzerland).

Based on the R^2^, the full model explained 62% of the variation in the number of compensated sheep per year in each NUTS3 region. A model including only the fixed terms (predator and prey abundance, landscape structure and the historical large carnivore presence) explained instead 13% of the variation, leaving the remaining 49% to the random part (hypothesis 6).

## Discussion

Our analysis revealed a wide variation with respect to all the components of the depredation and compensation process. Large carnivore densities, husbandry practices, protection measures, compensation systems, timing of coexistence with large carnivores, etc., all varied among, and within, the European countries considered in the study. Compensation systems mainly exhibited a country-to-country variation, with the exception of the Italian case in which the issue is managed at the regional level (Boitani et al., 2010). All the other variables considered, though, varied widely among the different NUTS3 units within the same country. In particular, husbandry practices and the use of livestock protection measures, which can have a strong effect on the reduction of large carnivore impact (Eklund, López-Bao, Tourani, Chapron, & Frank, 2017), did not exhibit a consistent pattern in most of the countries (see Table S4 and Appendix 1), but varied from region to region, likely as the result of a combination of environmental, social and historical processes, and because of the complexity of their implementation. Such multi-scale spatial variation is at the core of the challenges that human-large carnivore coexistence faces (Linnell, 2015): large carnivore populations are inherently trans-boundary and need a transboundary approach to their management (Linnell & Boitani, 2012), but most of the factors that determine the magnitude of their impact on human activities are influenced by local factors and require a local approach to be fully understood (van Eeden et al., 2018). This also highlights a partial limitation of our continental approach to the study of large carnivore depredation, as some information on the relevant factors in the depredation process were simply not available at the appropriate local scale and for the appropriate geographic extension required. Such limitations are revealed by the fact that the fixed part of our depredation model, in which the explicit variables were included, explained only 13% of the total variation in reported depredation levels. Our research approach, though, was not focused on explaining local variation, as on testing multiple broad scale hypotheses. When trying to reveal the effect of one or a few factors on the depredation process, the local scale is usually the most suitable, because it allows to gather high resolution data in a rather homogeneous geographic context (Eklund, López-Bao, Tourani, Chapron, & Frank, 2017). On the contrary, a large-scale approach is required when trying to assess the relative role of several components on the resulting large carnivore impact. A wider approach assured the necessary co-variation of all the components at a wider geographic scale, thus allowing to answer more general questions. This came at the cost of a coarser data resolution, but allowed us to produce answers to all our six research questions.

The first prediction we were able to test regarded the link between large carnivore distribution, their abundance and the resulting damage on livestock, an issue that is crucial impact mitigation. The debate about large carnivore impact often focuses on the questions of how many carnivores occur in a certain area, if they should be numerically reduced, and, if so, how many should be culled. On this and similar issues, the debate is usually highly polarized, under the implicit assumption that numbers are crucial when it comes to large carnivore damage (Treves, Krofel, & McManus, 2016). Although we were not able to directly test the effect of large carnivore abundance on impact, distribution proved to be a strong and reliable proxy, allowing us to extrapolate our conclusions with a certain level of confidence. To this regard, our results provide a nuanced answer to the question. In the case of wolves, and looking at the macroscopic continental gradient, a larger distribution (and likely higher abundance) implied higher levels of reported depredation; on the other hand, the link between large carnivore distribution and damage was weak and not significant for the other three large carnivore species, although the model suggested a positive relationship for them, too. Bautista et al. (2019) also found contrasting evidence of the link between large carnivore numbers and compensated damage. They revealed a positive relationship between the rate of range change in the last five decades and the costs for damage compensation in brown bears, but not in wolves and lynx (Bautista et al., 2019). These results suggest that distribution and abundance cannot be disregarded as irrelevant factors in livestock damage, and that management actions aimed at influencing them should be evaluated as an option, because they can affect damage. On the other hand, distribution and abundance alone are likely to be poor and weak predictors of large carnivore impact. Our analytical framework shows that a few carnivores can produce high levels of damage, when the totality of the environmental, historical, social and economic system favours it, whereas large populations can produce a very limited material impact, when the same components of the system reduce the probability that depredations occur.

Norway and Sweden, for example, share similar habitat and climatic conditions (although rather different landscape and terrain structures) and they are both experiencing an expansion of large carnivore ranges and numbers during recent decades, after a long period of absence or drastic reduction (Chapron et al., 2014). They display large differences, though, when it comes to the prevalent sheep husbandry practices and to the characteristics of their damage compensation systems. Sheep in Norway are traditionally free-ranging and unguarded on summer pastures and do not gather in flocks, whereas in Sweden the vast majority of them are raised in fenced fields all year round (Linnell & Cretois, 2020). Also, in Sweden the vast majority of compensation claims are based on a field inspection by state inspectors and only verified depredations are compensated, whereas in Norway only about 5-10% of damage compensations stem from a field inspection of a carcass, whereas the remaining 90-95% refers to payments made for missing animals which are assumed to be killed by large carnivores (Swenson & Andrén, 2005). Likely as a result of these social and administrative differences, Norway exhibited four times more compensated sheep heads than it would be expected based on large carnivore abundance in the country, whereas in Sweden compensation levels were about six times lower than expected by large carnivore abundance (Fig. 4).

A similar example of how relevant the anthropogenic component of the depredation process can be is provided by the Croatian results. Croatia hosts about 1,000 bears and 200 wolves, which overlap with about 400,000 sheep (Tab. 1). While there are by far more bears than wolves in the country, bear impact on livestock is close to zero (Majić, Marino Taussig de Bodonia, Huber, & Bunnefeld, 2011), whereas about 1,700 sheep are compensated each year as killed by wolves (Majić & Bath, 2010). A partial explanation for such differences lies in the fact that bears are omnivorous and feed on many other sources besides livestock, while wolves rely almost entirely on meat for their diet. Moreover, bears only partially overlap with the distribution of sheep farming areas in the country. Still, other components need to be considered. Bears are traditionally managed as a de facto game species in Croatia and the maintenance of a large population secures income for hunters in rural areas (Knott et al., 2014). Moreover, bear damage to sheep (and to beehives) is paid by local hunting associations, which are willing to pay the costs of compensation as a way to gain social acceptance for bear presence in the country (Majić et al., 2011). The whole system, which benefits from a traditional human-large carnivore relationship based on hunting and management at the local level, seems to be both socially and economically sustainable. On the other hand, wolves in Croatia are not a game species and therefore not perceived as a recreational or economic resource for hunters. Rather, they are seen mainly as human competitors both for livestock and for game, with social conflict being especially high in recently re-colonized areas (Majić & Bath, 2010). In this sense, the wolf damage compensation system in Croatia is similar to the ones commonly found in most European countries: compensation is managed at the national level and livestock losses are refunded after a field inspection, but farmers are often unsatisfied with the amount of the compensation and the long transaction times (Kaczensky at al., 2012). Overall, the number of wolf-related compensation payments in Croatia is several times higher than it would be expected based on wolf population size in the country, whereas bear damage is much lower than predicted by bear abundance (Fig. 4). Such differences in depredation patterns between two large carnivore species within the same country also highlight that solutions to human-large carnivore coexistence issues are bound to be species-specific, and that no recipes are valid for all contexts and all species. While comparative studies are useful to reveal patterns, actions and policies should be grounded in each local context and finely tuned for each large carnivore species. The good news resulting from our analysis of large carnivore depredation in Europe is that time seems to play in favour of a progressive reduction in the costs associated with human-large carnivore coexistence. Despite the potentially confounding effect of the unaccounted factors, our model provides a clear indication that longer periods of exposure are associated with a reduced impact of large carnivores on livestock. It is likely that the factor variable we used as a proxy for sympatry times was strongly correlated with a set of other variables, such as the level of human guarding of flocks, the use of livestock guarding dogs and electric fences, the choice of appropriate flock size, etc., which have been shown to reduce depredation levels in local studies (Eklund et al., 2017). Therefore, from a general point of view we could expect that time will allow the re-establishment of the appropriate co-adaptation tools (*sensu* Carter & Linnell 2016), which in turn will favour a reduction of the costs associated with sharing space with large carnivores in multiuse landscapes. However, there may well be more challenges with restoring traditional grazing practices with their associated protection measures in areas where they have been lost, as compared to maintaining them in areas where their use has been continuous. Moreover, the entire livestock industry is slowly changing due to social and economic drivers, which are causing the gradual abandonment of pastoral lifestyles (Linnell & Cretois, 2020). Without the appropriate management of the issues related to large carnivore impact on livestock husbandry, time may actually correspond to a progressive disappearance of small livestock breeding. This trend is further facilitated by the rules provided for by the Common Agricultural Policy (CAP) that has been applied in EU countries, and which tend to favour holdings with large numbers of heads, by definition more difficult to manage in a compatible way with the presence of predators. Finally, large carnivore populations are still expanding in most of the European countries (Chapron et al., 2014), making the economic sustainability of the whole compensation model unsure. Other models, such as risk-based or insurance-based compensation, are being tested, with contradictory results about their effectiveness and social acceptance (Marino, Braschi, Ricci, Salvatori, & Ciucci, 2016). The other relevant issue is that social conflict is often poorly related to material impact (Linnell, 2013). So, while technical tools and the appropriate mitigation policies might decrease the material impact of large carnivore presence on human livelihoods, the socio-cultural context may still generate conflict within and between stakeholders, unless careful attention is paid to governance structures (Linnell 2013a). Therefore, responsible agencies should try and focus their attention both on compensation and co-adaptation. While the reduction of large carnivore impact is a fundamental pre-requisite for the establishment of a sustainable long-term coexistence, there is also an urgent need for those participatory actions that consider the socio-cultural component of the process (Redpath et al., 2013) and that are more likely to increase the speed of the human-large carnivore re-adaptation process, thus progressively moving from an armed co-occurrence to a sustainable coexistence.

## Authors’ Contributions

V. Gervasi, O. Gimenez, J. Linnell and L. Boitani conceived the ideas and designed methodology; All authors contributed to data collection; V. Gervasi and O. Gimenez analysed the data; V. Gervasi, O. Gimenez and J. Linnell led the writing of the manuscript. All authors contributed critically to the drafts and gave final approval for publication.

## Supporting Information

Additional Supporting Information may be found in the online version of this article:

**Tab. S1** – Data sources for depredation data.

**Tab. S2** – Data sources for sheep distribution data.

**Tab. S3** – Data sources for large carnivore abundance data.

**Table S4** - Summary of the prevalent husbandry practices, damage reduction tools and compensation systems in the 10 European countries included in the large carnivore impact analysis, years 2010-2015.

**Appendix 1** – Description of the prevalent sheep husbandry practices, damage reduction systems and compensation systems in each of the 10 European countries included in the review and analysis of large carnivore damage compensation, years 2010-2015.

